# Genome Enrichment of Rare, Unknown Species from Complicated Microbiome by Nanopore Selective Sequencing

**DOI:** 10.1101/2022.02.13.480078

**Authors:** Yuhong Sun, Xiang Li, Qing Yang, Bixi Zhao, Ziqi Wu, Yu Xia

## Abstract

Rare species are vital members of a microbial community, but retrieving their genomes is difficult due to their low abundance. The ReadUntil (RU) approach allows nanopore devices to sequence specific DNA molecules selectively in real-time, which provides an opportunity for enriching rare species. However, there is still a gap in RU-based enriching of rare and unknown species in environmental samples whose community composition is unclear, and many species lack corresponding reference in public databases. Here we present metaRUpore to overcome this challenge. We applied metaRUpore to a thermophilic anaerobic digester (TAD) community, it successfully redirected the sequencing throughput from high-abundance populations to rare species while facilitating the recovery of 41 high-quality metagenome-assembled genomes (MAGs) at low sequencing effort. The simplicity and robustness of the approach make it accessible for labs with moderate computational resources and hold the potential to become the standard practice in future metagenomic sequencing of complicated microbiomes.

## 1 Introduction

Microbial communities are composed of a high number of rare species^1^. Rare species play a vital role in ecosystem health and stability^2^. For example, the slow-growing autotrophic microbes of ammonia-oxidizing bacteria or archaea (AOB/AOA) and anammox enable the rate-limiting step for natural nitrogen turnover^3,4^. Therefore, identifying the functional capacities of these rare species is essential to understanding the community dynamics and ecological function of a natural microbiome^2,3^.

The recovery of draft genomes (referred to as metagenome-assembled genomes, MAGs) from high-throughput metagenomic whole-genome sequencing (thereafter short as metagenomic) datasets ushered in a new era for understanding the ecological and evolutionary traits of the unculturable majority of natural microbiomes. However, high-quality (HQ, usually defined as >90% completeness with <5% contamination and the intact rRNA operon^44^) MAGs recovery for low abundant species is always difficult. In metagenomic sequencing, the low-abundance microorganisms are often missed or simply neglected due to low sequencing coverage. To get sufficient genome coverage of low-abundance species, extremely deep sequencing will be required. It would be a great waste if the study aims were to focus on rare species. Things can become more intractable during the data analyses that recovering the unknown genomes from hundreds of gigabytes to terabytes of data is a massive computational challenge^4^.

To raise coverage of rare taxa from a high-abundance background, molecular biology-based methods including hybrid capture or CRISPR-Cas9 enrichment are adapted in library preparation to enrich target^5,6^. On the other hand, depletion of high abundance species may serve the same purpose. Saponin-based host DNA depletion in human metagenomic communities is used for rapid clinical diagnosis of relatively low abundance pathogenic bacteria^7^. What is evident, however, is that these approaches require the use of extra reagents and preparatory procedures. This is compounded by the fact that they require known information about the enrichment or depletion targets in order to design the experiment, which does not appear to work for enriching low abundance species in metagenomic communities with unknown compositions.

Unlike the endeavors made prior to sequencing, Nanopore sequencing (Oxford Nanopore Technology, ONT) users can program their system to reverse the voltage polarity of the sequencing pore to eject reads identified as not of interest, which provides a potential solution to enrich for rare species in metagenomic samples. This ‘selective sequencing’ or Read Until (RU) strategy was first implemented by Loose and colleagues in 2016^8^. The earliest adopted dynamic time warp (DTW) algorithm-based approach could not scale to references larger than millions of bases, which limits its widespread usage^8^. With the similar goal of mapping streaming raw signal to DNA reference, UNCALLED has a lighter computational footprint than DTW^9^. Still, it requires abundant computational resources. The newly designed Readfish toolkit eliminates the need for complex signal mapping algorithms, and exploits existing ONT tools to provide a robust toolkit for designing and controlling selective sequencing experiments^10^. Until now, the application of RU is principally limited to the elimination of known host species^9, 10, 11^ or the enrichment of known targets such as mitogenomes of blood-feeding insects^12, 13^.

By ejecting dominant species while accepting low-abundance species, selective sequencing provides a potential solution to enrich rare species in metagenomic samples. Nonetheless, enrichment for low abundance species in real metagenomic samples by selective sequencing remains challenging because the community composition is never known, and a large proportion of the species lacks a corresponding reference in public databases. To specifically address such metagenomic-issue and to realize effective targeted enrichment of rare species within a complicated environment microbiome, here we introduced metaRUpore, a protocol consisting of know-how for configuring selective nanopore sequencing and necessary bioinformatic scripts to achieve efficient enrichment of rare species within a complicated environment microbiome. We initially assessed the efficacy of enriching low abundance species in a mock community. Based on this evaluation, we elaborated the principles and processes of metaRUpore and applied it to a thermophilic anaerobic digester (TAD) community that was treating waste sludge of a domestic wastewater treatment plant (WWTP). Meanwhile, we demonstrate a robust and effective procedure for assembling and binning HQ-MAGs from RU-based nanopore datasets. And an archaeal HQ-MAG retrieved from the TAD community revealed a giant (112Kbp) function-related genomic island, extending the evolutionary traits of the important *Bathyarchaeota* phylum.

## 2 Results

### *H. mediterrane* enrichment in a mock community

To evaluate nanopore performance on enriching low abundance species with RU, we firstly constructed a mock community. The *Haloferax mediterranei* strain which accounts for 1% of the mock community, was the target of our enrichment, while the other seven bacteria species were targets to be depleted during the RU run. In the mock run, a MinION flow cell was configured into two parts, where the first half of the channels did selective sequencing, and the other half did normal sequencing as a control. In the RU channels, the reads were basecalled and then mapped to a 33-M reference which contained all these eight microorganisms when they are being sequenced. A DNA molecule would be firstly sequenced for 0.4s before the obtained sequence was aligned to decide it should be sequenced continually or ejected. The average length of rejected reads was 537 bases, it demonstrated that the entire process of basecalling, mapping, and rejection decision could be completed in about 1.3s, based on the average nanopore sequencing seed of 400bp/s with R9.4.1 chemistry^10^. In the RU-delivered dataset, >99.9% of archaeal reads were kept while >99% of bacterial reads were ejected. *H. mediterranei* got enriched to the absolute dominant population within the community with a relative abundance of 62% in kept reads (Fig. 1a) with the coverage increased twice to 21.19× in RU data (Fig. 1b).

**Fig. 1.**
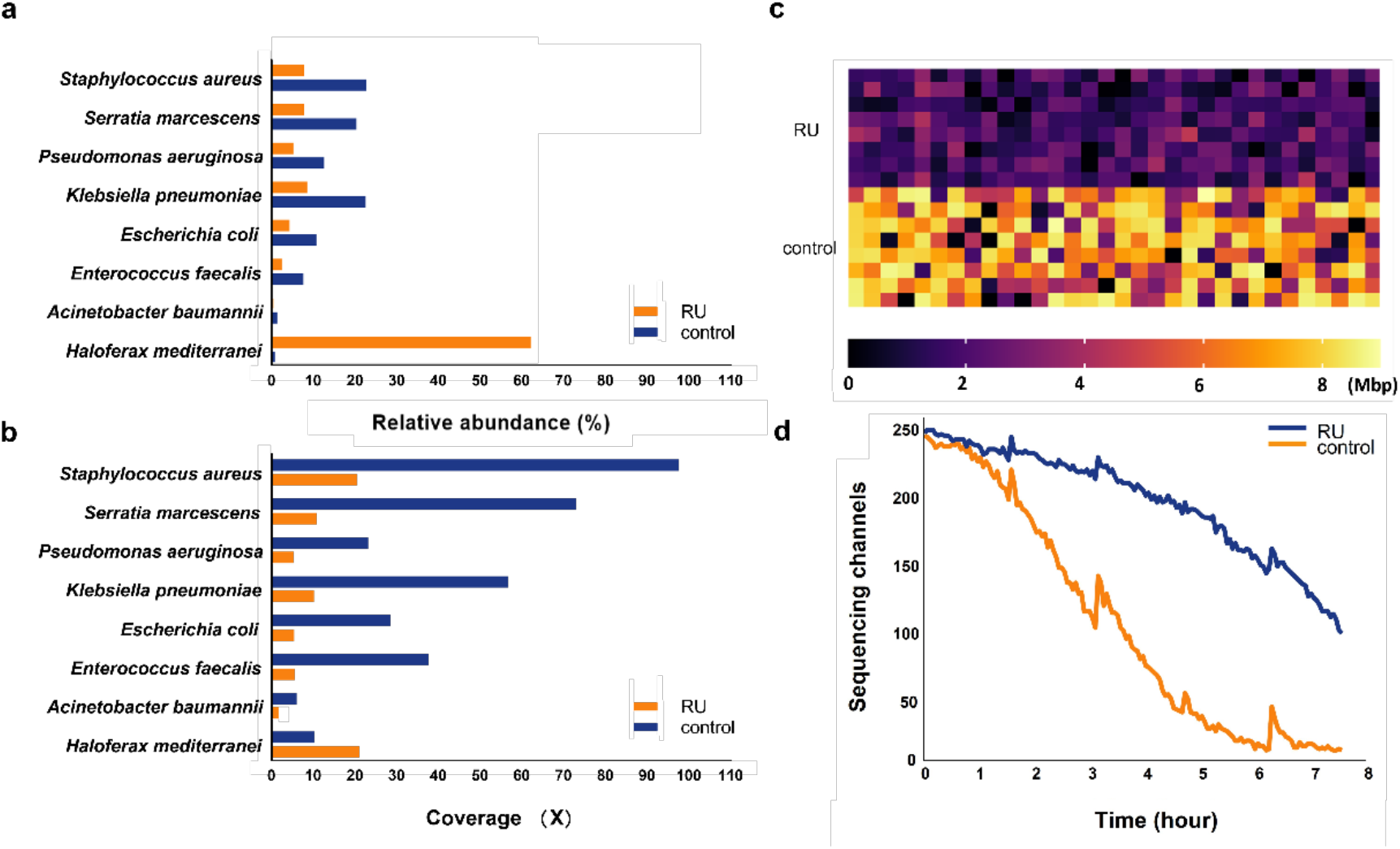
Enriching low abundance species in mock community with RU. **a**, Bar plot of the abundance of the seven microbial species in RU and control runs. **b**, Bar plot of the coverage of the seven microbial species’ genome in RU and control runs. **c**, heatmap of data yield per channel in RU and control runs, and **d**, plot of the number of sequencing channels over the course of the sequencing run.

Despite the high rejection precision and fairly ideal enrichment result, it must be noted that the total yield of selective sequencing was approximately 60% lower than that of normal sequencing (Fig. 1c). This reduction in throughput can be partly attributed to the increased idle time of each nanopore caused by a large number of ejections^9^. At an enriched target prevalence of 1% within a community, each nanopore ejected an average of 2,430 short fragments while 267 continuous long fragments were sequenced in a 7-hour run. In addition, a rapid drop in active channels happed after 1-hour sequencing in RU channels (Fig. 1 d and Supplementary Fig. 1) and the effective pore got depleted after 6-hour runtime which was 4 times shorter than normal run whose pores could normally last for 24 hours (Fig. 1d). Consequently, it’s critical to establish an appropriate target proportion for selective sequencing to achieve the best tradeoff between enrichment effectiveness and throughput loss. Fortunately, increasing sequencing effort could easily compensate for the RU-induced per flow cell throughput loss.

### In situ Metagenomic selective sequencing protocol and performance

We introduced a pipeline, MetaRUpore (https://github.com/sustc-xylab/metaRUpore), to selectively sequence rare populations in complex microbiome samples. The protocol consists of three consecutive steps (Fig. 2a): (1) 1h normal sequencing to obtain an overall picture of the community structure and the genomic profile of the dominant populations, (2) bioinformatics analysis to determine the reference and target dataset for optimized RU configuration, and (3) finally a 40h selective sequencing for enriching rare populations in the sample. The pore control of the nanopore device was implemented by Readfish^10^ which combines Guppy with minimap2^14^ to determine the eject/keep action for a pore.

**Fig. 2.**
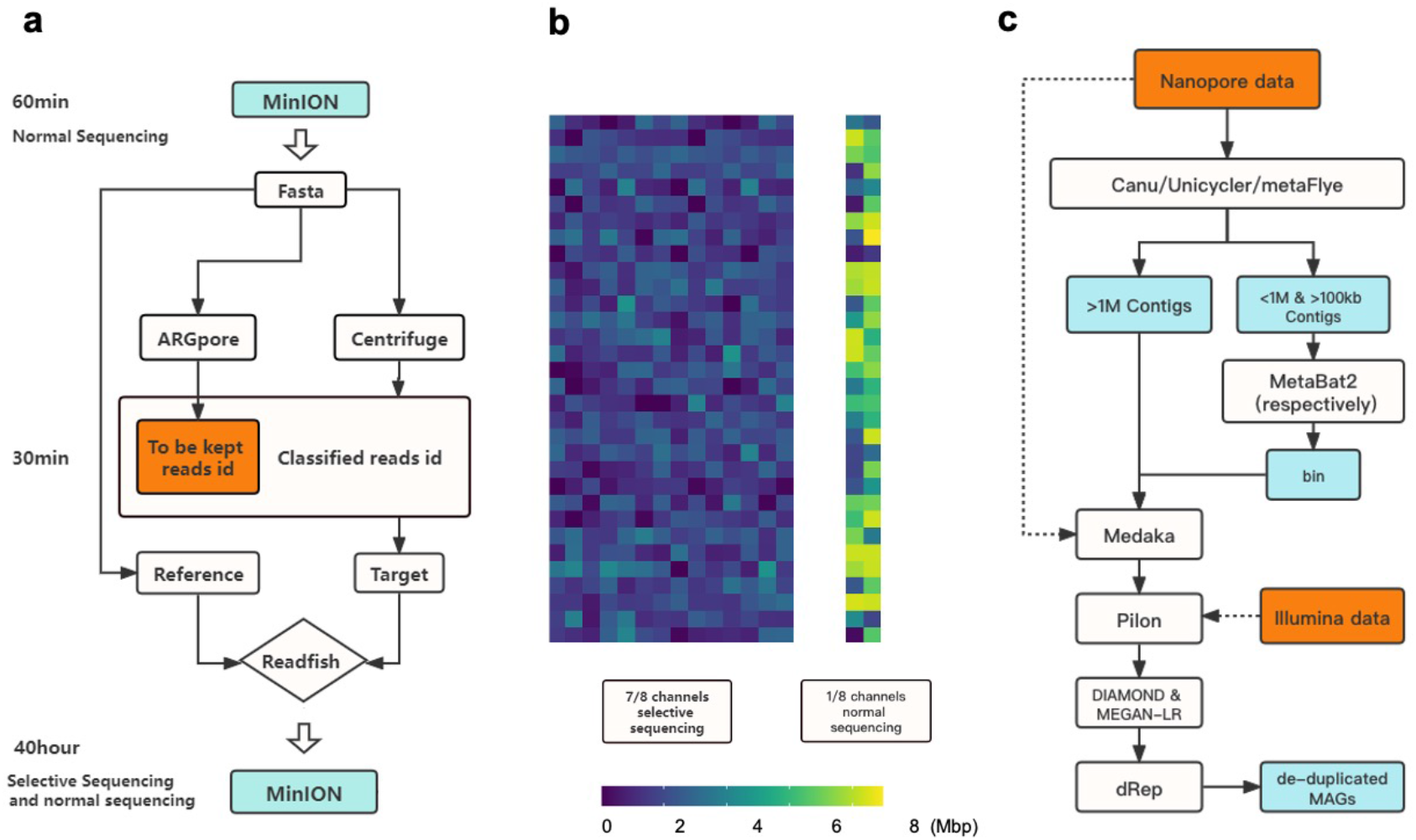
**a**, The workflow of metaRUpore. **b**, A MinION flow cell in metaRUpore is configured into two parts, 1/8th of the channels for normal sequencing and the remaining channels for selective sequencing. **c**, The bioinformatic workflow for HQ-MAGs retrieval based on datasets derived from nanopore selective sequencing and Illumina sequencing.

Here we show our results in applying the metaRUpore protocol to facilitate the genome recovery of rare populations within the TAD community, which consists of 2,977 OTUs with a Shannon index of 8.74, representing a typical diversity level of bioreactor systems (Supplementary Fig. 2). Rarefaction analysis demonstrated that the reads sequenced in the first 1 h normal sequencing already cover 90% of the overall diversity in the TAD community (Supplementary Fig. 5). Among the 125,606 reads sequenced, 66% of them could be assigned to a known reference by Centrifuge^15^. All of these classified reads obtained in the first 1 h run were set as the target for ejection in subsequent RU run as it mostly consisted of the known and abundant populations within the community. Notably, using whole-genome sequences from close species (same family or genus) as the reference for RU run will result in poor performance in ejecting the dominant populations because environmental microbiomes typically contain a high proportion of genetic fragments that are distinct from all the sequences deposited in whole-genome collections. In fact, even with the entire bacterial whole genome collection set as the ejection target, only an ejection efficiency of 22% was achieved in RU sequencing of the TAD community, leaving the community profile largely unchanged after selective sequencing. Another thing to note is that the classified reads obtained in the firstly 1h normal sequencing, inevitably contain genomic fragments from the rare and unknown populations we intend to enrich, which will result in incomplete genome coverage of rare populations in the sequences obtained in the RU channels. Therefore, a small fraction of the channels still needed to be set to normal sequencing in the subsequent 40h RU run and the delivered dataset needs to be assembled together with the RU-derived datasets. For our RU-sequencing of the TAD community, we set 1/8 channels to normal sequencing (--channels 1 448) (Fig. 2b). Our subsequent data analysis revealed that 29 HQ-MAGs would be missed if reads derived from selective sequencing were assembled alone. To further manipulate the selection, the users can manually select which taxa to keep during subsequent RU run; reads belonging to these taxa will be subtracted from the target dataset based on their taxonomic affiliations determined by ARGpore2^16^. For example, in our TAD community, we intended to keep all the archaea reads, so we eliminated them from the ejection target datasets. The entire aforementioned bioinformatic analysis can be completed in less than 30 min, such short suspension will not affect the flow cell chemistry and the subsequent RU run may directly start without refreshing the sequencing library.

The 40h RU run on one flow cell delivered 6.84 Gbp of effective long reads with an average read length of 3.46 kbp, while the normal sequencing channels produced 1.71 Gbp reads with an average read length of 3.60 kbp (Supplementary Fig. 3). To ensure adequate genome coverage, we sequenced the TAD community following metaRUpore protocol using three flow cells one by one on GridION X5. Given the concern to exhaust computation capacity on GridION X5, we did not test RU run with multiple flow cells sequenced simultaneously. RU sequencing using metaRUpore protocol resulted in a marked change in the community structure. As shown in the 3D density plot of phylogeny distribution of the overall TAD community (Fig. 3a), several density peaks of the original TAD community were depleted in the RU-run delivered datasets, indicating DNA of the high abundance populations of the TAD community was effectively ejected during RU-sequencing and the community got homogeneous with coverage of different populations become much more unified. Such unified coverage of different populations will help to minify the disparity of kmer frequency in the dataset, preventing kmers of the rare species from being filtered out as error-containing kmers due to coverage drop during the kmer-counting step of a *de novo* assembly algorithm^17, 18^.

**Fig. 3.**
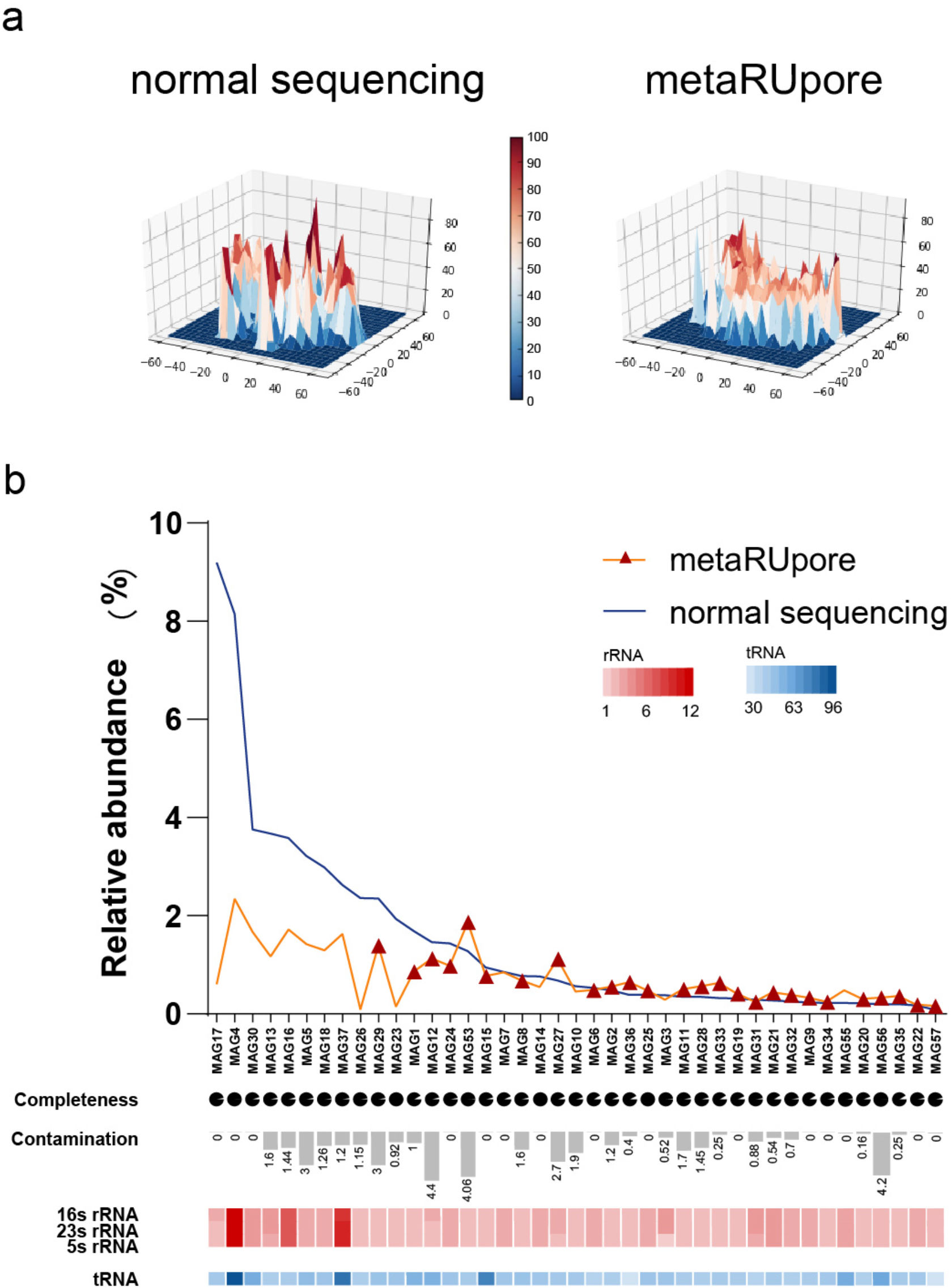
Performance of metaRUpore on recovery of high-quality MAGs in TAD community. **a**, 3D density plots of t-SNE downscaling results for normal sequencing datasets and selective sequencing datasets by metaRUpore at four base frequencies, showing that metaRUpore renders the community structure homogenous. **b**, The distribution of 41 retrieved HQ MAGs in normal and RU sequencing dataset. The red triangles indicate MAGs that were could only be assembled in the metaRUpore dataset. The pie chart and bar chart represent the level of genomic completeness and contamination by CheckM. The copy number of 16S rRNA, 23S rRNA, and 5S rRNA is represented by the red heatmap, while the copy number of tRNA is represented by the blue heatmap.

### Bioinformatics pipeline for *de novo* metagenomic assembly and genome recovery

As illustrated in the assembly pipeline (Fig. 2c), the 31G data consisting of RU and normal sequencing were assembled together respectively using three different assemblers, namely Canu^19^, Unicycler^20^, and metaFlye^21^. The basic statistics of assembled contigs were summarized in Supplementary Table 1. To improve the robustness of the binning, 139 > 1Mbp contigs were firstly picked, as the candidate of HQ genome^22^. The rest shorter contigs derived by the three assemblers were respectively binned by MetaBAT2^23^. Only contigs longer than 100 kbp were kept for subsequent binning. The MAGs retrieved above were subject to consensus correction by Medaka with nanopore data and polished by Pilon^24^ with Illumina short reads (SRs). Next, polished MAGs were further corrected for frame-shift errors using MEGAN-LR^22^ based on DIAMOND alignment against the *nr* database. Finally, MAGs obtained by the different assemblers were de-duplicated using dRep^25^ with a relatedness threshold of ANI > 0.95 to obtain species-level representative MAGs. Totally, we obtained 46 draft-quality MAGs after dereplication. Among them, 41 MAGs including 6 complete circular genomes were high-quality (HQ) (Supplementary Fig. 8 and Supplementary Table 2). 32 of these HQ MAGs were firstly picked single >1Mbp contigs, while the remaining 15 HQ MAGs were obtained by binning. All of these MAGs contained less than 13 contigs with an average N50 > 2 Mbp, demonstrating that they are highly continuous. In comparison, the normal nanopore sequencing dataset yielded 29 draft-quality MAGs, including 16 HQ MAGs. 15 of them were included in the 41 HQ MAGs retrieved by metaRUpore strategy (Supplementary Fig. 8). Worth noting is that the 26 HQ MAGs that are additionally obtained by RU-based selective sequencing were mainly from the rare populations of the TAD community (Fig. 3b). Additionally, evident coverage reduction was observed in the dominant populations that the coverage of MAG17, MAG4, and MAG30, which together accounted for 21% of the TAD community, dramatically reduced by 78% after RU-based selective sequencing (Fig. 3b and Supplementary Table 3), demonstrating the effectiveness of metaRUpore protocol in eliminating dominant populations during sequencing. Despite the lowered overall throughput, coverage of the rare species MAG33, MAG35, MAG57, and MAG56 was doubled at the current sequencing effort and the application of the metaRUpore protocol has reduced the abundance limit for HQ-MAG recovery in the TAD community to 0.7%. It could be expected that by using additional flow cells, HQ-MAGs could be obtained for populations with even lower prevalence.

## 3 Discussion

### Complete genomes recovered from TAD community

The 41 HQ MAGs introduce 5 new phyla, namely *WOR-3, OLB16, Omnitrophota, Gemmatimonadota*, and *Deferribacterota*, into the global HQ genome collection of AD microbiome^26^ (Fig. 4). Furthermore, our MAGs show much better integrity and continuity than those in the previous collection assembled with SRs in terms of N50, number of contigs as well as intact rRNA operon. Additionally, evolutional traits analysis reveals a much more conservative scale of gene flow based on HQ genomes we assembled than that based on fragmented MAGs^27^ (Supplementary Fig. 9) .

**Fig. 4.**
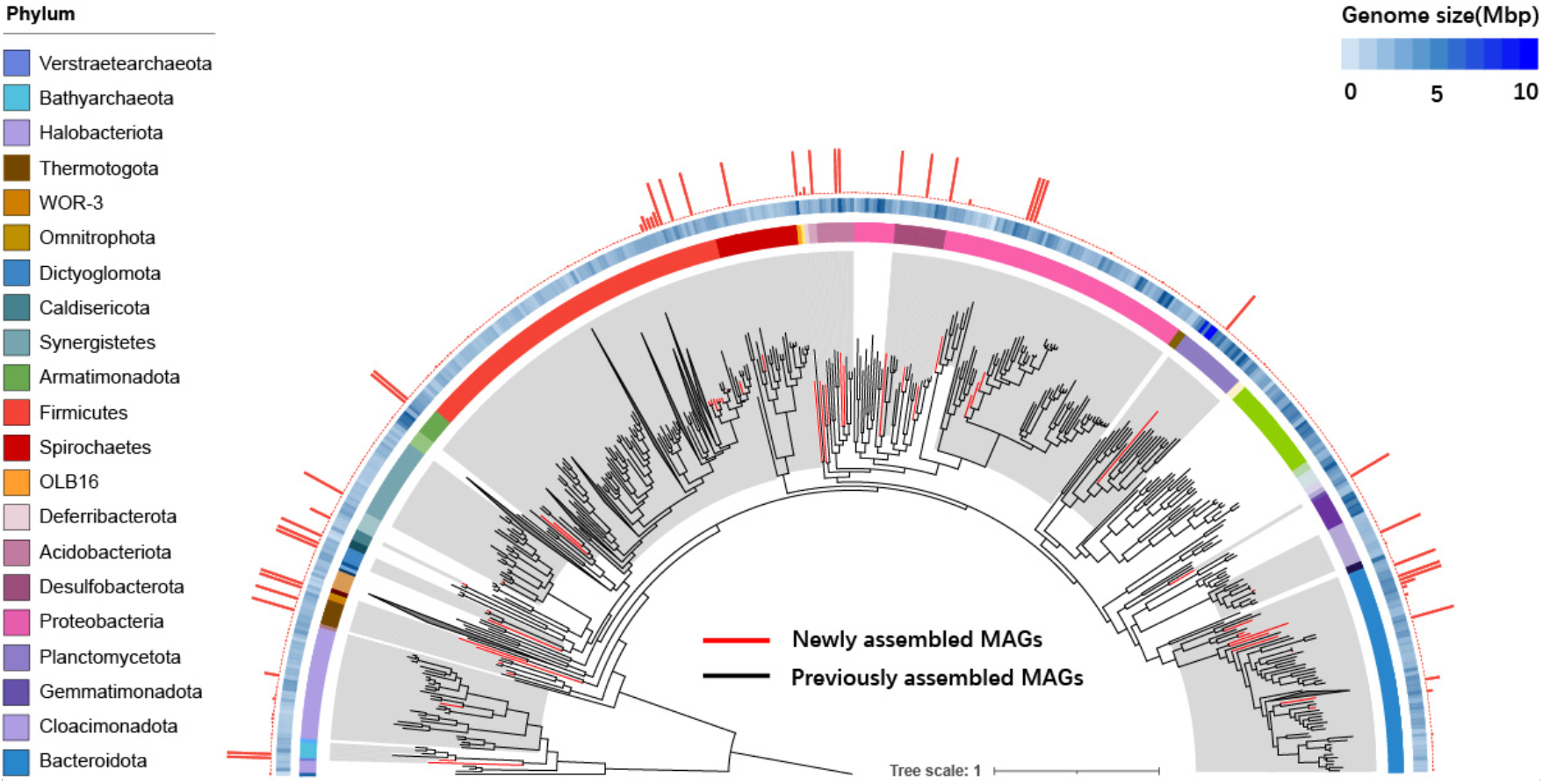
Phylogenomics of MAGs in anerobic reactor. A phylogenetic tree was constructed from 41 HQ-MAGs derived by metaRUpore (red branches) and 1,108 HQ-MAGs collection derived from other AD systems (black branches). External circles represent, respectively: (1) taxonomic assignment at phylum level, (2) genome size (heatmap), (3) bar plot representing the genome continuity, which is calculated as the reciprocal of the number of contigs. The grey shaded areas indicate phyla with near-complete genomes obtained by metaRUpore,and the name of each phylum is in the legend on the left.

### Versatile metabolic capacities of *Bathyarchaeota* phylum in TAD community

*Bathyarchaeota* was recently recognized as a methanogenesis contributor^28^ that may play active roles in global biogeochemical cycles^31^. However, the absence of pure cultures of the phyla has hampered our understanding of their ecological functions and evolutionary positions from a genome-centric perspective^29,30^. Genomes reported for this phylum so far are highly fragmented (Fig 5a). In this work, MetaRUpore has boosted the abundance of *Bathyarchaeota* in the TAD community from 0.19% to 0.32%, facilitating its genome recovery as MAG56, which to the best of our knowledge, is the first complete genome for this phylum. MAG56 represented a novel *Bathyarchaeota* lineage with the closest neighbor being Bathy-5 (Fig 5b). The genome size of MAG56 is 1.9Mbp, notably larger than the average size of previously assembled genomes of *Bathyarchaeota* phylum (1.23Mbp)^29,30,31^. *Bathyarchaeota* was previously proposed to have methyl-dependent hydrogenotrophic methanogenic potential^28,32^ as MAGs recovered from deep aquifers^34^ possess an MCR-like complex. However, no MCR homology could be detected in MAG56. Given the complete nature of the genome obtained in this study, a functioning methanogenic pathway in the TAD community lineage of *Bathyarchaeota* seemed implausible.

Remarkably, we found three genomic islands (GIs) (Fig 5a) in MAG56 with the largest being 36 kbp in length. These GIs were always missing in previously genomes assembled by short reads due to the defective resolving of repetitive fragments flanking the exogenous genetic island^33,34^. In the largest GIs of 36 Kbp, we identified six copies of Tyrosine recombinase (*xerA, xerC*, or *xerD*), which had previously been reported to facilitate the insertion of gene islands into the host chromosome by catalyzing site-specific, energy-independent DNA recombination^34,36^. Additionally, we identified a heat shock protein, *HtpX*, that may contribute to the heat shock response facilitating the cell’s survival in a thermophilic environment. Collectively, this GI represents a highly mobile fitness island^33^ that offers selective advantages for the archaeal population within the thermophilic digester community. And the recovery of complete MAGs by metaRUpore undoubtedly enabled the discovery of the role of large GIs in shaping *Bathyarchaeota*’s evolution.

**Fig. 5.**
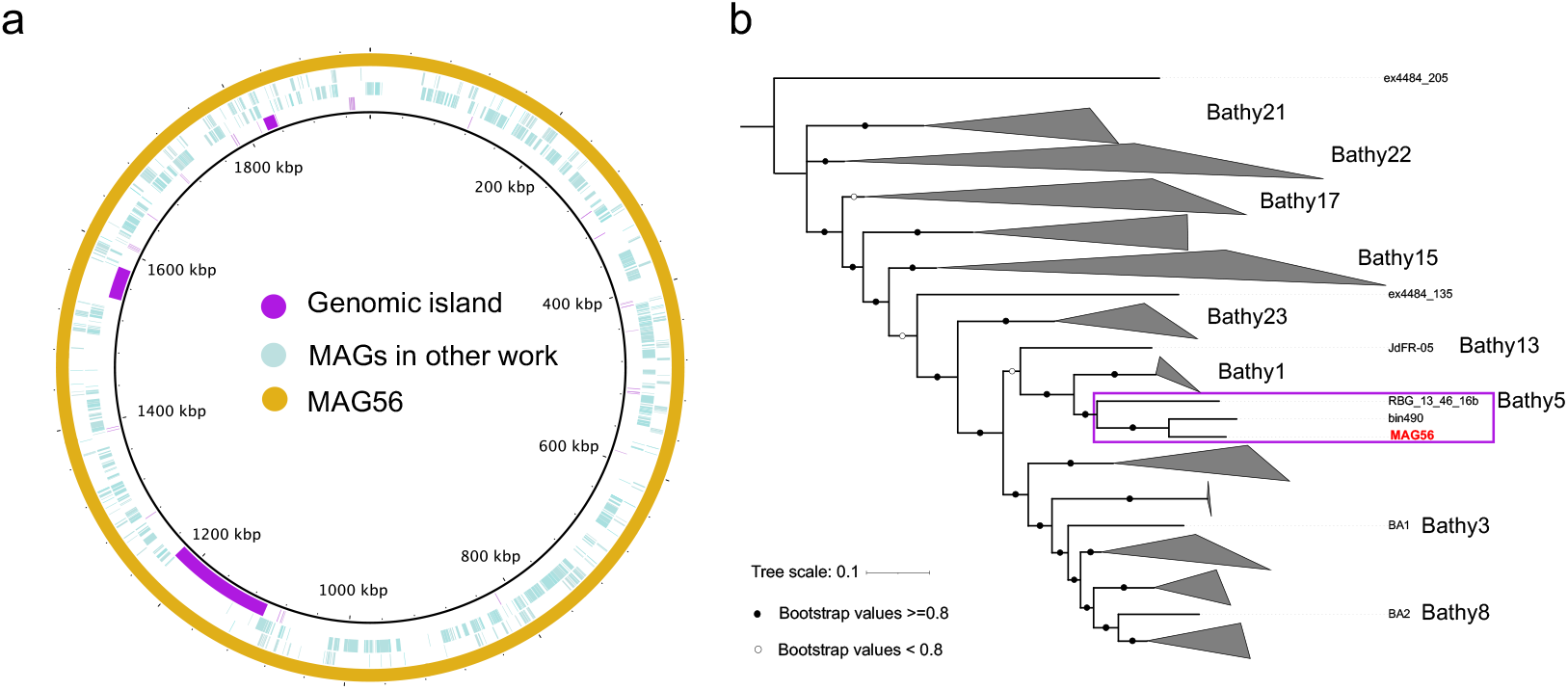
a, Genomes comparison of MAG56 and other MAGs of Bathyarchaeota from prior research. The outermost ring stands for the circular genome of MAG56 reconstructed by metaRUpore. The second to third circles from the outside represent the MAGs of phylum Bathyarchaeota reconstructed by short reads-only assembly method (MAGs covered by purple boxes in Figure 5c). The innermost purple circle represents the genomic island. b, A Maximum Likelihood Tree showing the phylogeny of Bathyarchaeota based on the MAGs from the current study (MAG56) and prior research^29^. Bootstrap values for these phylogenies are shown with open (< 80%) and filled (≥ 80%) circles.

Overall, we proposed metaRUpore, a method for enriching low-abundance and undiscovered microorganisms in complex microbial communities based on nanopore selective sequencing. The heuristic ejecting targets determined through initial short-term *de novo* sequencing of the dominant populations, overcome the constraints imposed by the absence of reference genomes for selective sequencing of complex communities. metaRUpore unifies the sequenced community structure and increases the genome coverage of low-abundance species, facilitating the assembly of additional HQ genomes of rare species within the microbiota. HQ MAGs retrieved from the TAD community by metaRUpore contribute to the building of a more comprehensive database of AD-associated microbes, which will ultimately allow for an in-depth understanding of their biological characteristics. More importantly, metaRUpore protocol is robust and requires minimal modification to the experimental procedure of nanopore library construction and sequencing, making it easily applicable to metagenomic investigations of other environmental microbiomes. Even though selective sequencing for the rare sphere is inevitably associated with a reduction in per-flow cell data yield. Future implementation of the RU API on PromethION will easily provide a throughput boost, overcoming the coverage barrier and enabling complete genome recovery of rare species with even lower abundance from complex microbiomes using the metaRUpore protocol.

## 4 Methods

### Sampling and DNA extraction

Genomic DNA of the eight microorganisms of the mock community was extracted by QIAamp DNA Micro Kit (50). Samples for TAD community were taken when the methanogenic bacteria were at their highest activity. Genomic DNA of the TAD community samples was extracted by QIAGEN DNeasyR PowerSoilR Kit (100). DNA concentration was determined using the Life Technologies Qubit high sensitivity assay kits. The quality of the DNA was measured by Thermo Scientific™ NanoDrop™ to assure that it all met the requirements for library construction.

### Construction of the synthetic mocks

We synthesized a mock community of eight microorganisms, of which Archaea accounted for 1% and the other seven bacteria species shared the rest equally based on DNA concentration determined from qubit average measurements. The archaeal species is *Haloferax mediterranei* and these seven bacteria are *Acinetobacter baumannii, Enterococcus faecalis, Escherichia coli, Klebsiella pneumoniae, Pseudomonas aeruginosa, Serratia marcescens*, and Staphylococcus aureus.

### Library construction and Sequencing

All sequencing libraries were constructed using the ONT Ligation Sequencing Kit (no. SQK-LSK109) according to the manufacturer’s instructions. When preparing the reactor sample libraries, in order to remove as many very short DNA fragments as possible, 0.4X beads was used for each step of the cleanup, and therefore the initial amount of genomic DNA was increased to 2ug to ensure a sufficient amount of DNA of the final library. ONT MinION flowcells v.R9.4.1 were used for all sequencing on an ONT GridION.

### Selective sequencing via metaRUpore

The execution of metaRUpore to enrich for unknown low abundance taxa is divided into the following three steps: firstly, a period (in this case 60 min) of normal sequencing is performed to generate reference file for selective sequncing using Readfish^10^ which should contain the vast majority of taxa in the community. Next, the sequenced data is fed into metaRUpore to obtain the reference and target needed to configure Readfish TOML for selective sequencing. During this time, it is advisable to keep the MinION flowcell with the DNA library in a 4°C refrigerator to avoid the loss of activity of the nanopores affecting the subsequent sequencing. We put the reference and target paths into the TOML file and set config_name = “dna_r9.4.1_450bps_fast”, single_on = unblock, multi_on = unblock, single_off = stop_receiving, multi_off = stop_receiving, no_seq = proceed, no_map = proceed. As recommended by the author of Readfish, we deactivated adapter scaling by editing the config files (dna_r9.4.1_450bps_fast.cfg) in the guppy data directory. Next, selective sequencing was started. the configuration on MinKNOW was the same as for normal sequencing. Readfish runs at the same time as the sequencing starts.

### Analysis of long-read sequence data

Sequencing-derived fastq reads were performed adaptor trimming using Porechop (<u>GitHub - rrwick/Porechop</u>) (version 0.2.2) with default settings. These reads were subsequently assembled by the three tools: Canu^19^ (version 2.2, default setting except -nanopore, genomeSize=3m, maxInputCoverage=10000, corOutCoverage=10000, corMhapSensitivity=high, corMinCoverage=0, redMemory=32, oeaMemory=32, batMemory=200 useGrid=false), Unicycler^20^ (version 0.4.9b, default setting except -t 40, --keep 3) and Flye^17^ (version 2.8.3, default setting except –nano-raw, -- threads 50, --plasmids, --meta, --debug). Generated contigs that was at least 1Mbp in length were regarded as potential whole-chromosome sequence. Among the remaining contigs that are less than 1Mbp, we did metagenomic binning for the contigs that are greater than 100kbp in length. Metabat2^21^ (version 2.12.1 with default setting) is used to respectively binning the contigs assembled by above three assemblers.

Next, we took multiple steps to correct the >1Mbp potential chromosome and bins we obtained. Firstly, we used nanopore data to perform consensus correction on them using Medaka (<u>GitHub - nanoporetech/medaka</u>)(version 1.4.3, default setting except -t 20, -m r941_min_high_g360). They were then further corrected with the short reads data using Pilon^23^ (version 1.24 with default setting except --fix all, --vcf). We used DIAMOND^35^ (version 0.9.24) to align the Pilon polished potential chromosome (with default settings except -f 100 -p 40 -v --log --long-reads -c1 -b12) against the NBCI–NR database^38^ (July 2021). We used daa-meganizer in MEGAN Community Edition suite^39^ (version 6.21.7, run with default settings except --longReads, --lcaAlgorithm longReads, -- lcaCoveragePercent 51, --readAssignmentMode alignedBases) to format the .daa output file and receive frame-shift corrected sequence with ‘Export Frame-Shift Corrected Reads’ option.

We checked the completeness and contamination of these potential genomes with CheckM^40^ (version v1.0.12, run with default setting except lineage_wf, -t 20). All the putative genomes were de-replicated using the dRep^25^ (version 3.2.2, run with default setting except -p 40 -sa 0.95 –genomeInfo) to get species-level unique MAGs. Next, gene annotations were obtained using Prokka^41^ (version 1.13). Microbial taxonomic classifications were assigned using GTDB-Tk^42^ (version 1.3.0, GTDB-Tk reference data version r89).

### Calculation of the abundance and assessment of the quality of MAG

Abundance was calculated from both selective sequencing data and normal sequencing data, by mapping these data to the MAGs using minimap2^14^ (version 2.17) separately using the following flags -ax map-ont -t 40. We used samtools^43^ (version 1.11) to extract .sam file that matched each MAG individually. The abundance of each MAG is calculated by dividing the number of bases in all reads in this .sam file by the total number of bases selectively sequenced or normally sequenced. Analogously, sorted .bam files were used in the calculation of coverage of the MAGs.

We defined high-quality (HQ) MAGs as encoding multiple rRNA genes (23S/16S/5S), SCG-completeness > 90% and contamination < 5%^44^. Draft-quality (DQ) MAGs means MAGs having > 70% SCG-completeness, < 10% contamination, and the presence of 16S rRNA. While if a MAG meets all of the DQ criteria but misses 16S rRNA were regarded as low-quality (LQ) genomes.

## Code availability

The metaRUpore workflow is available on the GitHub page: https://github.com/sustc-xylab/metaRUpore.

## Availability of data and materials

The raw nucleotide sequence data (both Illumina and Nanopore) used in the present study has been deposited in the NCBI database under project ID PRJNA794848.

## Acknowledgements

This work was supported by the National Key Research and Development Program of China (Grant No. 2021YFA1202500), the National Natural Science Foundation of China (Grant No. 42007216) and Shenzhen Science and Technology Innovation Committee (Grant No. JCYJ20210324104412033). Also, we want to thank the Center for Computational Science and Engineering at Southern University of Science and Technology (SUSTech) and core research facilities at SUSTech to provide quality resources and services.

## Conflict of interests

The authors claim no conflict of interests.

## Notes

https://github.com/sustc-xylab/metaRUpore.

### Competing Interest Statement

The authors have declared no competing interest.

