## Supplementary Information for "Genome Enrichment of Rare, Unknown Species from Complicated Microbiome by Nanopore Selective Sequencing"

### Supplementary Figures

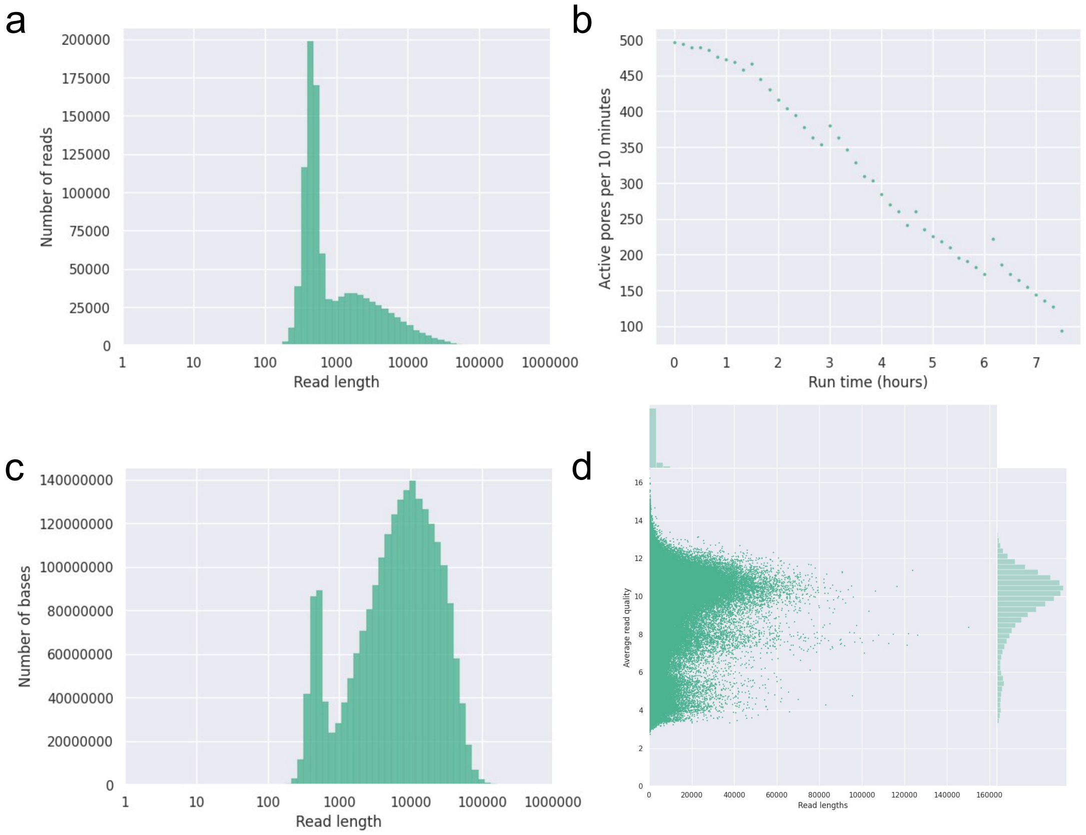


*Supplementary Figure 1: Selective sequencing report of mock community. a) Histogram of lengths after log transformation. b) Number of total active pores over time. c) Weighted histogram of read lengths after log transformation. d) Plot of read lengths versus average read quality.*

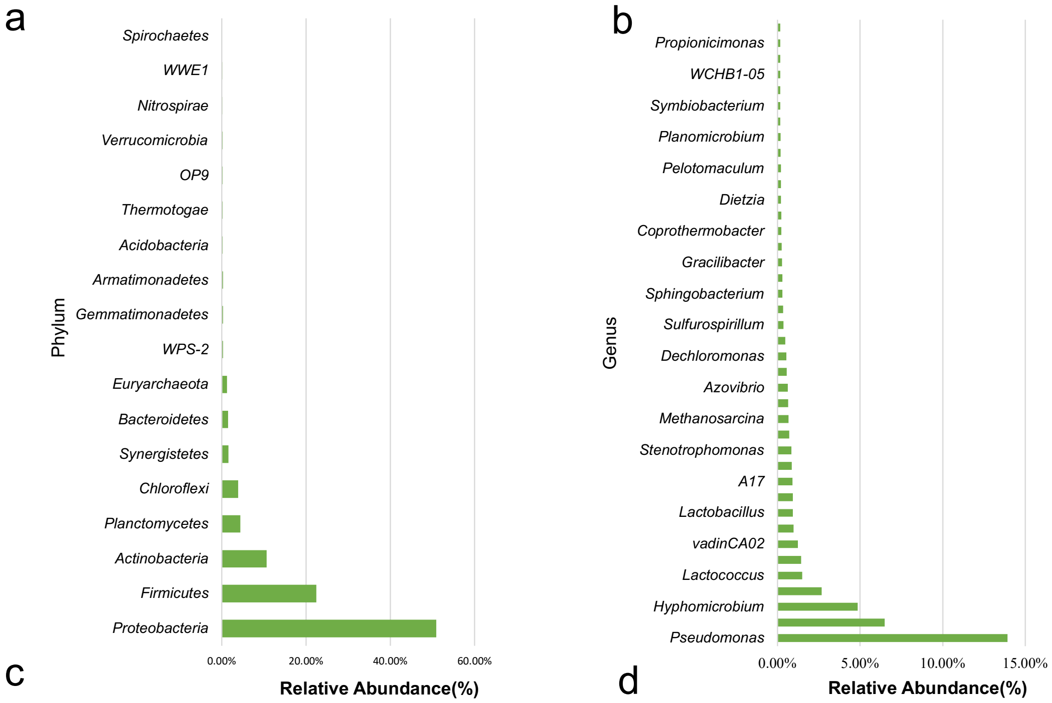


c

| | Phylum | Class | Order | Family | Genus |
|---|---|---|---|---|---|
| Classified Ratio | 99.73% | 99.03% | 94.34% | 78.96% | 46.80% |

d

| Sample | Chao1 | Reads | Shannon |
|---|---|---|---|
| TAD | 7545.83 | 237778 | 8.74 |

*Supplementary Figure 2: a) Phylum, b) Genus level community structure of the thermophilic anaerobic digester (TAD) community. c) Classified ratio of each taxonomy level. d) Alpha diversity index based on metagenome extracted 16S rRNA.*

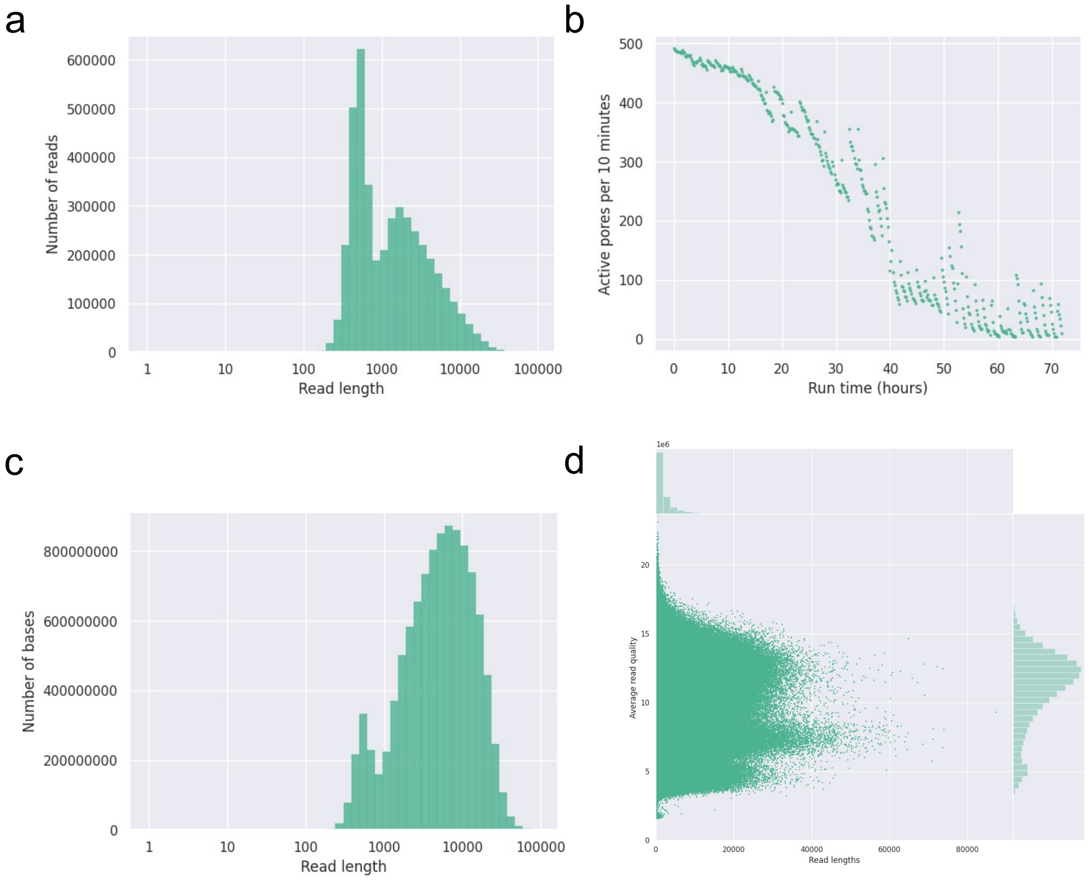


*Supplementary Figure 3: Selective sequencing report of TAD community. a) Histogram of lengths after log transformation. b) Number of total active pores over time. c) Weighted histogram of read lengths after log transformation. d) Plot of read lengths versus average read quality.*

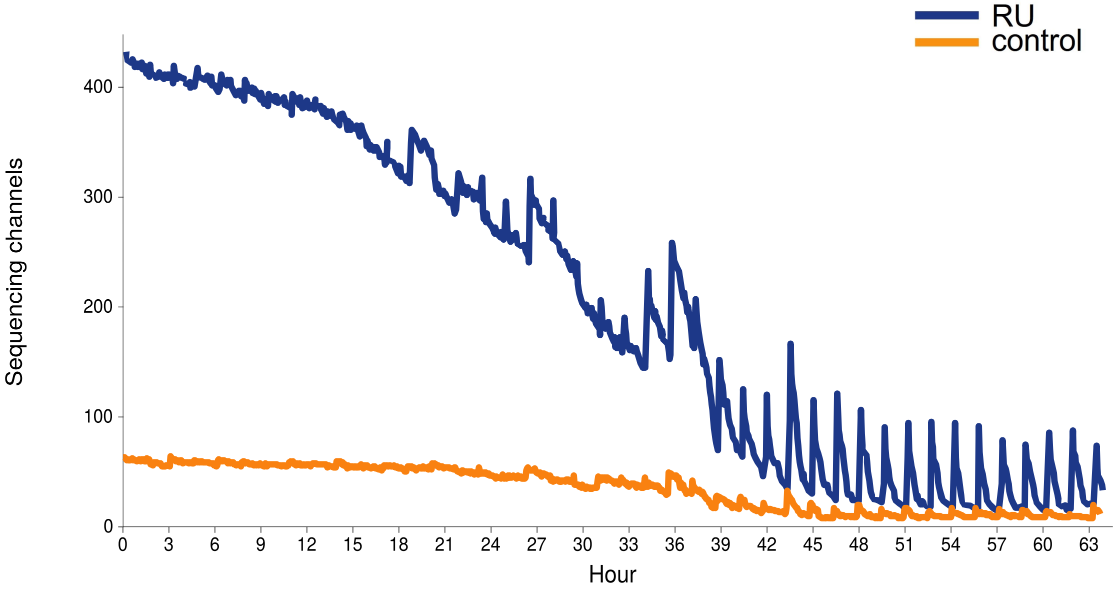


*Supplementary Figure 4: The number of sequencing channels over the course of the sequencing run in TAD community. It shows that active pore loss speed of RU-channels wasl faster than that of the control channels by the slop of the*

*line.*

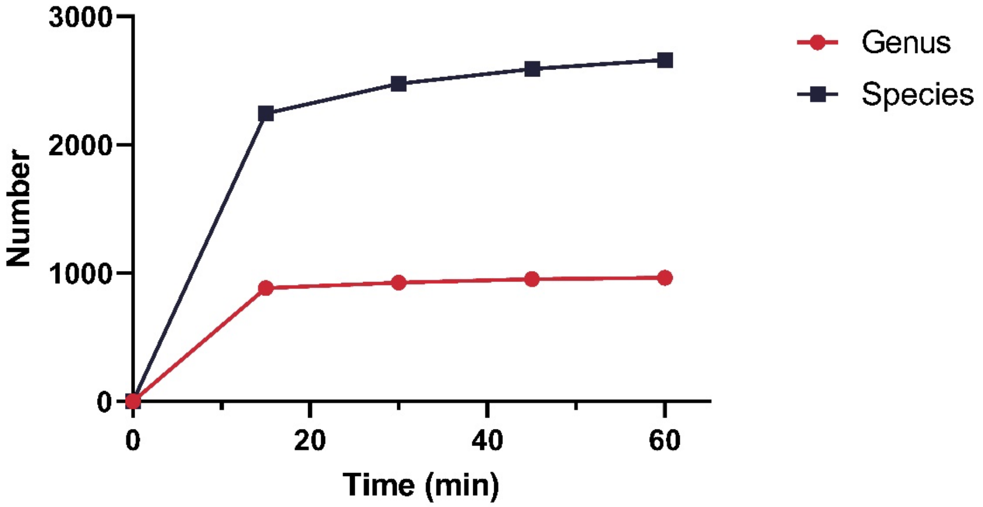


*Supplementary Figure 5: Rarefaction analysis of nanopore sequencing data. The Y-axis is the number of species or genus annotated to by Centrifuge. The curve is close to saturation at 60min.*

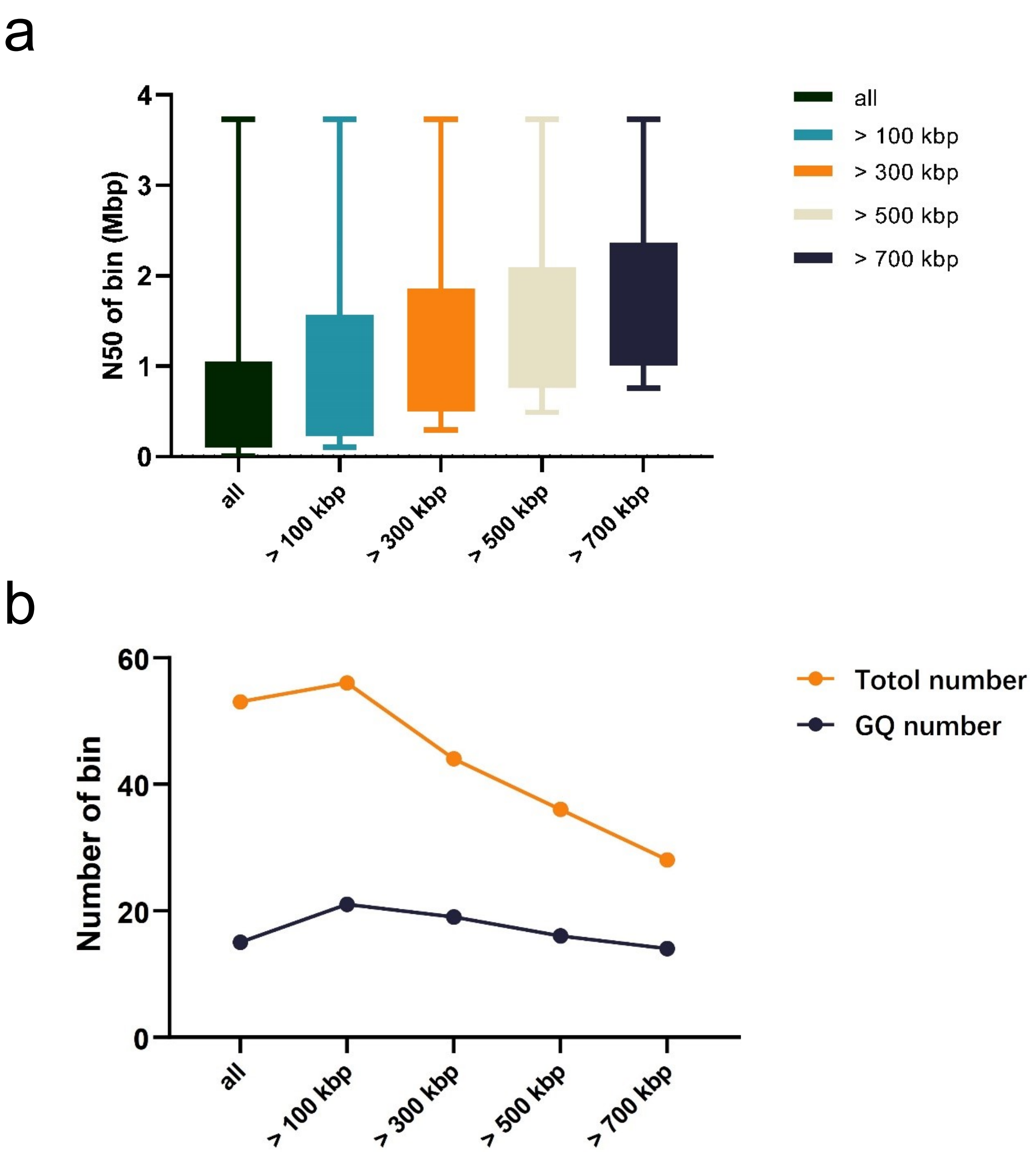


*Supplementary Figure 6: The quality and quantity of bins obtained for contigs of different lengths. We grouped the contigs <1M into five categories: >700 kbp, >500 kbp, >300 kbp, >100 kbp, and all contigs and binned them separately. As a result, binning with >100kb contigs could achieve the greatest balance between quantity and quality of MAGs, so we finally chose 100kbp as a tradeoff for binning. a) N50 of the bin obtained from contigs of different length groups. b) Number or good-quality number of the bin obtained from contigs of different length groups. Good quality bins mean they have > 80% completeness and < 5% contamination, with the potential to be corrected to high-quality bins.*

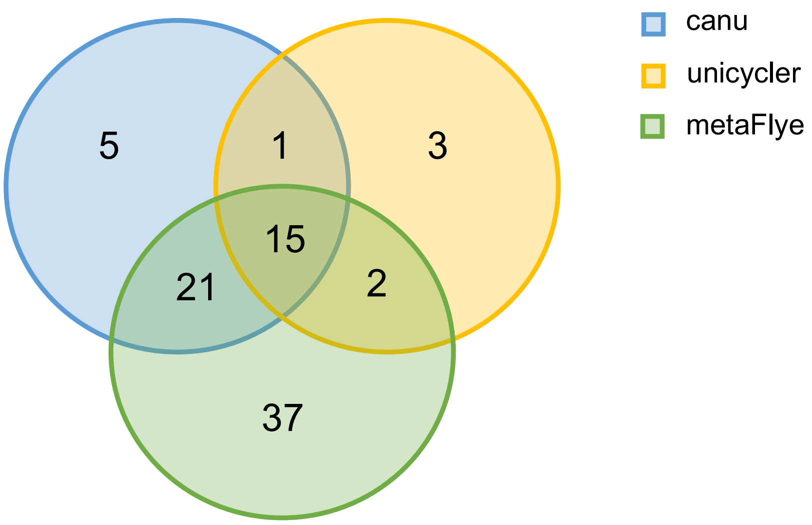


*Supplementary Figure 7: Venn diagram of the number of >1Mbp contigs assembled from canu, unicycler, and metaflye, respectively. We assembled the nanopore data with canu, unicycler, and metaflye, respectively, and de-duplicated them by dRep with a relatedness threshold of ANI > 0.95. We found that the three tools produced duplicate >1 Mbp contigs, but each tool was able to assemble additional contigs.*

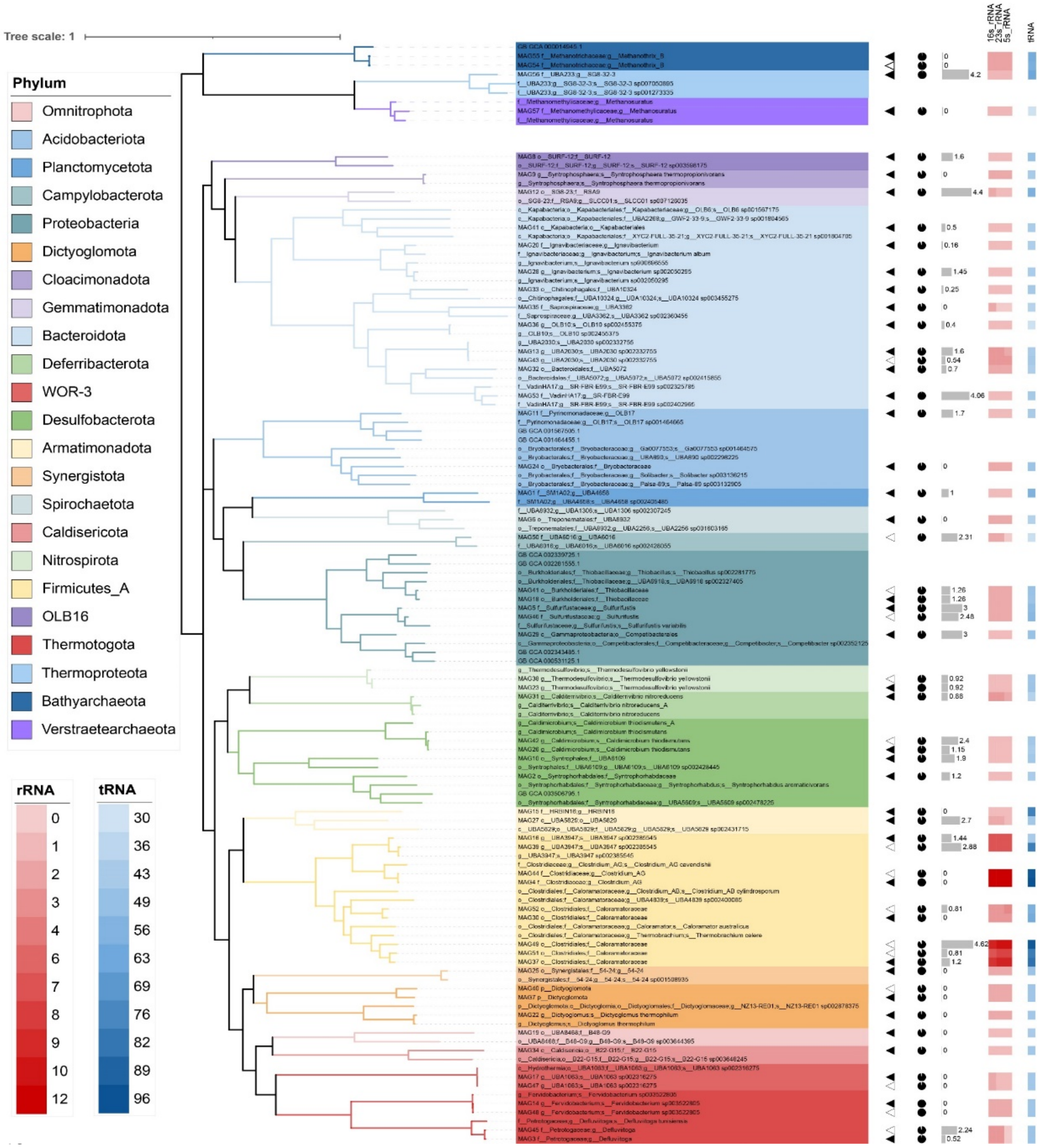


*Supplementary Figure 8: A phylogenetic tree was constructed from 57 HQ genomes derived from the TAD community and reference genomes. The solid triangles represent the 41 MAGs assembled from the metaRUpore dataset and the hollow triangles represent the 16 MAGs assembled from the normal sequencing dataset. The different coloured branches of the tree represent phyla, the pie chart represents genomic completeness and the bar chart represents genomic contamination. The copy number of 16S rRNA, 23S rRNA, and 5S rRNA is represented by the red heat map from left to right, while the copy number of tRNA is represented by the blue heat map.*

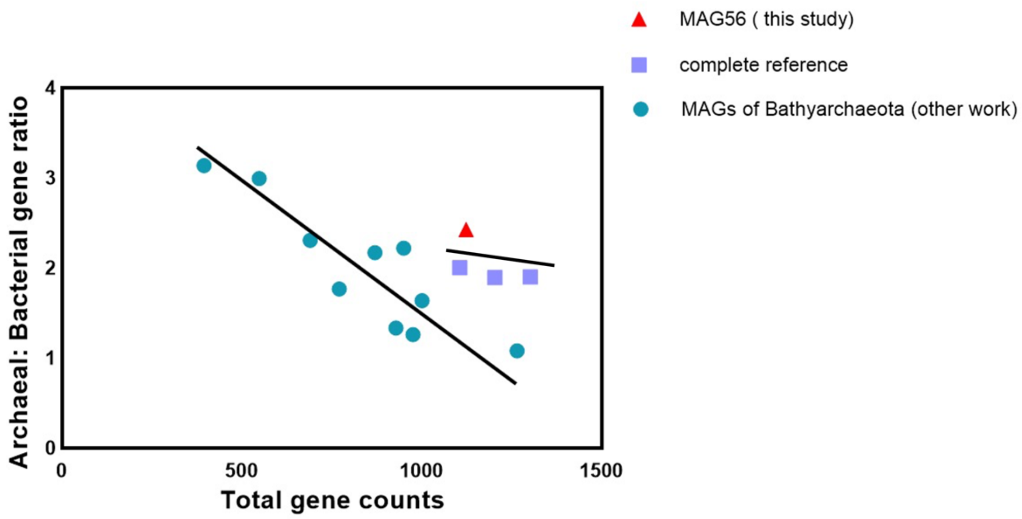


*Supplementary Figure 9: Correlation between the number of genes of each genome and the Archael: Bacterial gene ratio.*

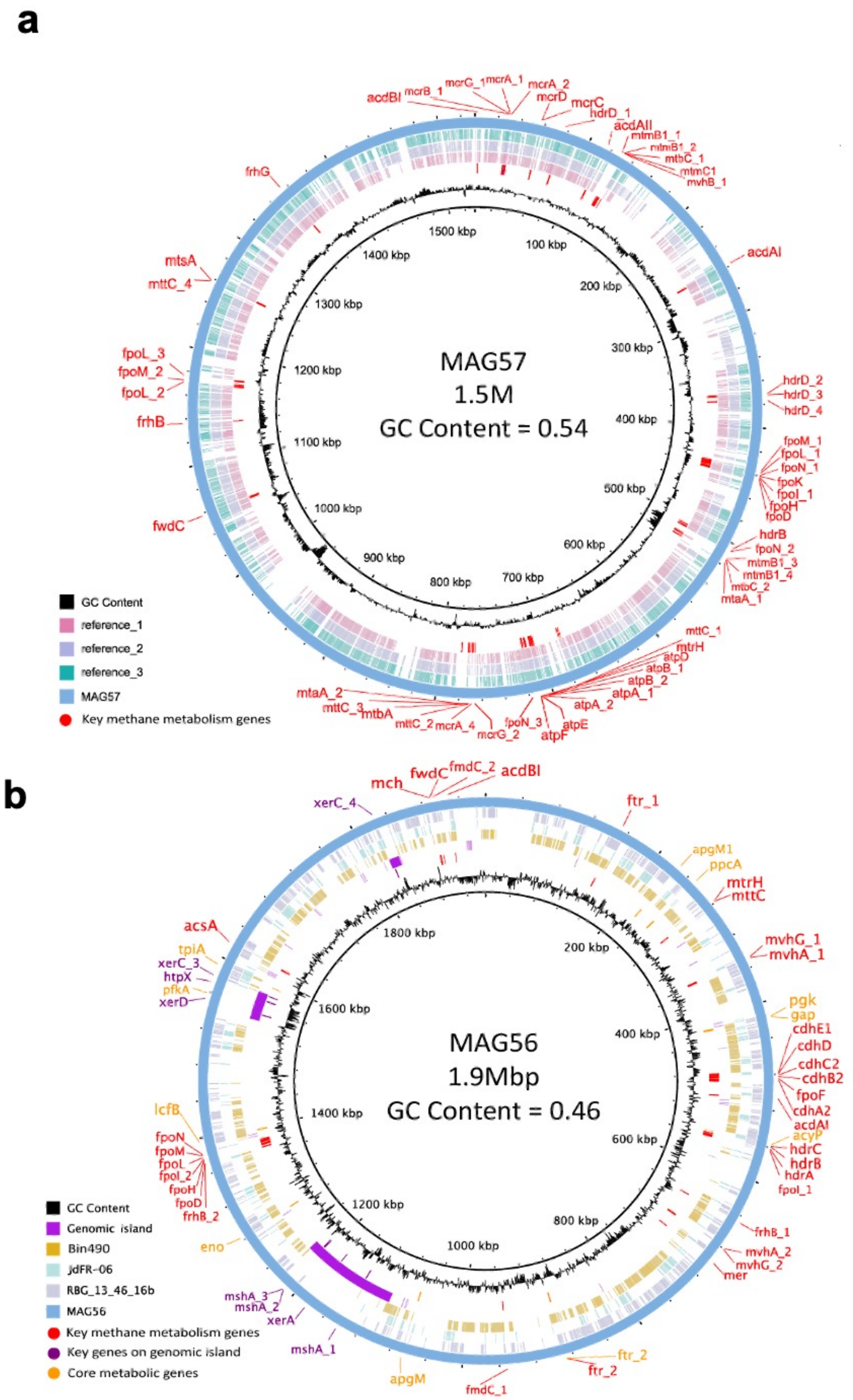


*Supplementary Figure 10: a) Genomes comparison of MAG57 and reference MAGs. The outermost ring stands for the circular genome of MAG57*

*reconstructed by metaRUpore. The second to fourth circles from the outside represent the MAGs of phylum Verstraetearchaeota reconstructed by short reads-only assembly method, which was mapped to MAG57. The two innermost circles from the outside to the inside indicated the key methane metabolism predicted genes by Prokka and GC content, respectively. b) Genomes comparison of MAG56 and reference MAGs. The outermost ring stands for the circular genome of MAG56 reconstructed by metaRUpore. The second to fourth circles from the outside represent the MAG, which was mapped to MAG56. The fifth purple circle represents the genomic island. The sixth circle from the outside indicated the key methane metabolism genes (red), key genes on a genomic island (purple) and core metabolic genes (orange) predicted by Prokka and the innermost circles represent GC content.*

### Additionally results and discussion

**Versatile metabolic capacities of Verstraetearchaeota and Bathyarchaeota phylum in TAD community.**

A complete genome of *Methanosauratus petracarbonis* affiliated with archaeal phylum Verstraetearchaeota was recovered as MAG57. The genome size of MAG57 is 1.5M and the GC content is 0.54. The abundance of *Methanosauratus petracarbonis* in TAD community was 0.075 %, which got doubled through selective sequencing, enabling successful retrieval of its entire genome.

MAG57 contains key genes for methane production (*mcrABG* and ancillary genes *mcrCD*)[4] as well as genes for methylamine utilization (*mtaA, mtbA, mtmBC, mtbBC, mttC, mtrH*). The reduction of heterodisulfide (CoM-SS-CoB) to ferredoxin could be accomplished by the coupling of exergonic H2-dependent heterodisulfide reductase (*hdrB*) and F420-non-reducing hydrogenase (*mvhB*). Meanwhile, the cytosolic complex of F420H2 dehydrogenases (*fpo*) consisted of consecutively located *fpoM, fpoL, fpoN, fpoK, fpoI, fpoH and fpoD*, can reoxidize the reduced ferredoxin while pumping protons across the cytoplasmic membrane to produce a proton gradient that drives the ATP synthesis via an archaeal-type ATP synthase. Additionally, *HdrD*, which is present in three copies in MAG57 and other Verstraetearchaeota genomes, may directly interact with the *fpo* complex and act as an energy-converting ferredoxin: heterodisulfide oxido-reductase. Furthermore, genes for hydrogenotrophic and acetoclastic methanogenesis pathways were absent in MAG57, a nearly complete genome of Methanosauratus petracarbonis species, consolidating the species' obligate H2-dependent methylotrophic methanogenesis capability[1,2]. Notably, while unusual for microorganisms involved in methane metabolism, the exit of adenosine diphosphate (ADP)-forming acetate synthetase (*Acd*) in MAG57 demonstrates that it can convert Acetyl-CoA to acetate, allowing for energy production via substrate-level phosphorylation[1]. Collectively, the coupling of obligate H2-dependent methylotrophic methanogensis and acetate-producing fermentative pathway of *Methanosauratus petracarbonis*'s genomic repertoire found in MAG57, reveals a unique ecological niche for carbon turnover and energy conservation in digestive systems rich of reduced methylated carbon compounds.

In this work, MetaRUpore has boosted the abundance of Bathyarchaeota in TAD community, facilitating its genome recovery as MAG56. MAG56 appeared to be capable of utilizing sugars as a carbon source and generating acetyl-CoA via the Embden–Meyerhof–Parnas (EMP) pathway (a nearly complete operon of *pfk, tpi, gap, pgk, apg, eno, ppc*) and pyruvate-ferredoxin oxidoreductase (*por*). ADP-forming acetyl-CoA synthase (*acd*) could then produce ATP and acetate, and this fermentative lifestyle was predicted to be the metabolic mode of several *mcr*-devoid Bathyarchaeota genomes[2,6] Besides that, MAG56 possessed key genes for the autotrophic reductive acetyl-CoA (Wood–Ljungdahl, WL) pathway (*fwd, ftr, mch, cdh*), implying its ability to utilize tetrahydromethanopterin (H4MPT) as the C1-carrier for autotrophic carbon fixation, which is an energy-generating process prevalent in archaea[5]. Additionally, the critical genes for lipid and benzoate degradation (*lcfB* and *acyP*) found in the MAG56 genome demonstrated its capacity to exploit lipid and benzoate as a source of carbon and energy. These core metabolic potentials of MAG56 are consistent with previous studies, consolidating Bathyarchaeota's organoautotrophic life strategy capable of utilizing a diverse array of carbon sources[3,5].
